# BayesENproteomics: Bayesian elastic nets for quantification of proteoforms in complex samples

**DOI:** 10.1101/295527

**Authors:** Venkatesh Mallikarjun, Stephen M. Richardson, Joe Swift

**Affiliations:** Wellcome Centre for Cell-Matrix Research; Division of Cell Matrix Biology and Regenerative Medicine, School of Biological Sciences, Faculty of Biology, Medicine and Health, Manchester Academic Health Science Centre, University of Manchester, M13 9PL, UK

**Keywords:** Label-free proteomics, post translational modification, Bayesian linear regression, primary cells and tissue, Markov chain Monte Carlo

## Abstract

Multivariate regression modelling provides a statistically powerful means of quantifying the effects of a given treatment while compensating for sources of variation and noise, such as variability between human donors and the behaviour of different peptides during mass spectrometry. However, methods to quantify endogenous post-translational modifications (PTMs) are typically reliant on summary statistical methods that fail to consider sources of variability such as changes in levels of the parent protein. Here, we compare three multivariate regression methods, including a novel Bayesian elastic net algorithm (BayesENproteomics) that enables assessment of relative protein abundances while also quantifying identified PTMs for each protein. We tested the ability of these methods to accurately quantify expression of proteins in a mixed-species benchmark experiment, and to quantify synthetic PTMs induced by stable isotope labelling. Finally, we extended our regression pipeline to calculate fold changes at the pathway level, providing a complement to commonly used enrichment analysis. Our results show that BayesENproteomics can quantify changes to protein levels across a broad dynamic range while also accurately quantifying PTM and pathway-level fold changes. Raw data has been deposited to the ProteomeXchange with identifiers PXD012784, PXD012782 and PXD012772. BayesENproteomics is available for Matlab: www.github.com/VenkMallikarjun/BayesENproteomics and Python3: www.github.com/VenkMallikarjun/BENPPy

## Introduction

Spiraling costs of drug/therapeutic development and a low probability of success have driven an increase in the use of models based on patient-derived material in an attempt to determine translational potential prior to clinical studies. However, unlike samples from genetically homogeneous, completely inbred model organisms, patient samples can possess substantial differences between individuals. This variability between samples can make discerning a statistically significant effect extremely difficult using the standard statistical methods commonly used in biology. The lack of statistical power caused by poor signal-to-noise ratios in primary samples and the difficulty in obtaining large donor cohorts can compromise potentially promising biological studies, resulting in wasted time and prohibitive expenditure.

The problem of high inter-individual differences is rarely more acute than in high-throughput omics experiments where subtle differences in the levels of individual features within samples (e.g. transcripts, peptides derived from proteins, etc.) are compounded over many such features, demonstrating quantifiable inter-individual differences even between genetically identical twins (Brodin et al., 2015). This is especially relevant to typical mass spectrometry (MS) proteomic methods wherein quantities associated with multiple features (i.e. peptides) are used in the calculation of a fold change for a single protein.

In bottom-up MS-based proteomics, proteins are enzymatically digested into peptides and the abundances of these individual peptides are used to derive protein quantifications. A core assumption is that peptide abundance is proportional to protein abundance. However, each peptide has different physico-chemical properties that mean they each behave differently during sample preparation or within the mass spectrometer itself. For instance, residues surrounding cleavage sites can affect efficiency of enzymatic digestion (Lawless and Hubbard, 2012; Rodriguez et al., 2008); some peptides also ionise less efficiently than others (Abaye et al., 2011) and in data-dependent analysis, peptides of similar mass-to-charge (m/z) ratios may compete during co-elution or ionisation (Schliekelman and Liu, 2014), potentially biasing the peptides finally detected. These discrepancies can be further exacerbated by biologically relevant post-translational modifications (PTMs) that alter the behaviour of individual peptides, resulting in substantial differences in measured intensities for peptides that belong to the same protein, even in purified protein samples. Even with normalisation, differences in the behaviour of individual features between donors and experimental treatments are difficult to account for.

It was recently shown that combining MS datasets from multiple fractions and extraction conditions could achieve near-complete coverage of the human cell proteome, including >14000 protein isoforms (Bekker-Jensen et al., 2017). Interestingly, the same study also identified ~10000 phosphorylation sites and ~7000 acetylation sites without specific enrichment. New sample preparation techniques and the increasing sensitivity of MS instrumentation have increased coverage of PTMs. However, with this increase in sensitivity comes the problem of resolving differential regulation of different forms of the same protein (proteoforms), as not all peptides belonging to a given protein respond in the same way to a given treatment, an assumption made by many commonly used quantification methods.

Common statistical methods for dealing with variability include the summarisation of peptide intensities using averages, sums or medians to give subject-level protein quantification (Goeminne et al., 2015). However, these subject-level summary models fail to account for differences in behaviour of peptides belonging to the same protein during analysis, and thus are prone to perturbation by outlier peptides. They are also highly dependent on obtaining large numbers of peptides from each protein and thus can suffer from low statistical power when analysing less extreme fold changes in small or low abundance proteins (e.g. transcription factors). To address this, previous statistical models have focused on exclusion of outlier peptides to improve protein quantification (Forshed, 2013; Swift et al., 2013a), at the cost of a loss of information relating to different proteoforms for a given protein; or have attempted to deduce differences in the abundance of peptides belonging to different sub-populations of the same protein (Henao et al., 2013, 2012; Webb-Robertson et al., 2014; Zeng et al., 2017).

Multivariate regression has been shown to be a powerful means of analysing complex omic experiments (Smyth, 2006). For proteomics specifically, linear regression models represent a powerful means of teasing apart different, known sources of variability in peptide intensities, with the aim of returning the variability caused by a given experimental condition. Linear modelling has become the standard for transcriptomic analysis thanks to the widely used LIMMA package (Smyth, 2004) and has been successfully adapted for MS data analysis. For example, in the peptide-based model proposed by Choi et al. (2014) with donor effects added:

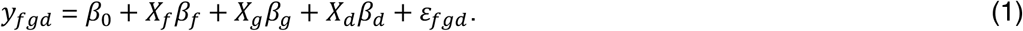

Where, for a given protein, *y*_*fgd*_ (the **response variable**) corresponds to the observed log_2_ intensity for peptide *f* of that protein under experimental condition *g* from donor *d*. **Predictor variables** *β*_*f*_, *β*_*g*_ and *β*_*d*_ correspond to the effect sizes of peptide *f*, group *g* and donor *d*, respectively. *β*_*0*_ represents the intercept term. *ε* corresponds to a Gaussian error term centred on 0 with a variance (*σ*^2^) specific to each protein. The purpose of the regression calculation was to find how much the slope of the resulting line differed for a given experimental treatment (i.e. the effect size *β*_*g*_ caused by the experimental treatment of interest *g*). In this instance the design matrix ***X*** consisted solely of categorical variables encoded as binary variables of size *n* × *p* (where *n* = number of observations and *p* = number of *β*s).

However, peptide-based linear models can be prone to overfitting, wherein predictive power is lost as the model attempts to fit the noise within the data rather than any overall trends. This may be due to the effects of outlier peptides, such as those that possess a biologically relevant PTM with a fold change different to that of its parent protein, or simply due to peptide mis-identification. The effects of outlier peptides on the final linear model fit can be mediated by various weighting strategies (e.g. see Goeminne et al. (2016)). However, for modelling changes in PTM abundance, it would be useful to obtain effect sizes for peptides that specifically interact with a given treatment and separate them from effect sizes for other parameters that may be interacting with these peptides. Linear modelling is more flexible than the name suggests and can also handle interactions between predictor variables; for example, where an experimental condition has an effect on the intensity of an individual peptide – independent of its effect on the protein as a whole – possibly due to some biologically important PTM of that specific peptide. For example, we can extend the model in (1) to allow separation of peptide:treatment and peptide:donor effects in studies using primary human samples, as in (2) (see also the model in Clough *et al*, 2009).

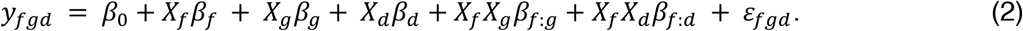

Wherein *y*_*fgd*_ corresponds to the intensity of peptide *f*, in experimental group *g*, from protein *i*, extracted from technical replicate *r* of donor *d*. The *β*s correspond to the fitted coefficients for each predictor variable. Note the interactions coefficients *β*_*f*:*g*_, and *β*_*f*:*d*_ in (2), denoting variability in individual peptide intensities, *f*, caused by the experimental treatment *g* or largely unknown features of the individual donor *d*. From here on we refer to (1) as linear and (2) as non-linear.

Unfortunately, overfitting becomes particularly acute in models that attempt to deal with many potential sources of variability, some of which may interact or correlate with one-another, and where the number of parameters to be estimated exceeds the number of observations. The inclusion of interaction terms in a model has potential to provide important biological insight regarding different isoforms and PTMs of a given protein (collectively called proteoforms) – vital mechanisms of post-translational regulation of cellular function. However, interaction terms may not be necessary for all proteins in a dataset, and the decision as to whether or not to include them would need to be made on a protein-by-protein basis. Importantly, while protein-based models are simpler and less prone to overfitting (because they do not include *β*_*f*_ or related interaction terms), peptide-based regression models have been shown to provide greater statistical power (Clough et al., 2009; Goeminne et al., 2015). Furthermore, by definition only peptide-based models are capable of providing quantifications for PTMs or differential splicing as variability in peptide behaviour is lost during protein-level summarisation. As we are interesting simultaneously quantifying protein and PTM fold changes, this eliminates protein-based models from consideration.

Simpler models (e.g. those used by (Choi et al. (2014) and (Goeminne et al. (2016)) minimise the potential for overfitting found in more complex models. However, when many potential sources of variability exist it may be necessary to fit increasingly complex models wherein the choice of terms included, particularly any interaction terms, may need to be decided on a protein-by-protein basis. Failure to select the correct terms could result in models failing to fit correctly, or even at all, especially if complex interaction terms are included when not needed. However, not including these terms may mean that subtle differences in peptide behaviour that occur only in specific proteins may go unaccounted for, resulting in a loss of accuracy. Thus, a means of automatically selecting appropriate features from a more complex model for use in fitting a final, sparse model is necessary.

Regularisation can be employed to minimise overfitting and perform feature selection. Regularisation involves attaching penalty weights to each coefficient (*β*), minimising its contribution to the final fit. The nature of the penalty can differ depending on the type of regularisation used. In the ridge regression algorithm used by Goeminne et al. (2016) an “L_2_” penalty is applied wherein *β*s are shrunk according to their squared norm, minimising the contribution of less important coefficients in explaining the total model variance. Although ridge regression shrinks coefficient estimates down to emphasise the most important *β*s, all *β*s remain non-zero. Another form of regularisation is LASSO (least absolute shrinkage and selection operator), which shrinks *β*s according to their “L_1_” absolute norm. With LASSO, some *β*s can be set to exactly zero, excluding them from the model and giving rise to so-called “sparse” solutions. Several variations of these methods have been proposed, including an “elastic net” variation that combines properties of both ridge regression and LASSO to perform both model selection and shrinkage of remaining non-zero *β*s (Zou and Hastie, 2005). These frequentist regularisation methods have Bayesian parallels depending on the prior distributions the *β* posteriors are sampled from, with L_2_ regularisation being equivalent to sampling of *β*s from a Gaussian distribution, L_1_ being equivalent to sampling from a Laplace (double exponential) prior and the elastic net being equivalent to *β* sampling from an intermediate distribution (Bornn et al., 2010).

Here we develop a Bayesian elastic net algorithm to provide regularised protein-specific models (BayesENproteomics), considering potential donor variability and interactions between specific peptides and experimental treatments and/or donor effects. This has the advantage in implicitly allowing for any individual peptide to behave differently from its identified protein group in response to treatment or donor effects. This means that BayesENproteomics does not assume that peptides have been identified correctly – a point we enforce by incorporating observation weights based on peptide identification confidence. However, unlike previous Bayesian regression models that attempt to accommodate variable peptide behaviour within a protein (Henao et al., 2013, 2012; Webb-Robertson et al., 2014), BayesENproteomics can still provide fold change estimates for protein groups containing peptides with specific identifications, maintaining interpretability of subsequent results by presenting relative quantification of the “dominant” proteoform for that protein (separating different isoforms if unique peptides are available), whilst also presenting values for differentially abundant PTMs for further investigation. We benchmark BayesENproteomics against other commonly used peptide-centric regression methods. Finally, we extend our linear modelling pipeline to encompass pathway analysis as a complement to standard enrichment analysis. This enables complete analysis of a dataset from MS1 intensities to functional interpretation of observed proteoform changes.

## Results

### BayesENproteomics shows increased sensitivity and accuracy in estimating protein fold changes compared to other regression models

To produce a benchmark dataset with known (ground truth) fold changes, peptides prepared from primary human mesenchymal stem cell (MSC) lysates and female C57BL/6J mouse skin were mixed in ratios of 3:1, 1:1 and 1:3 and analysed with a Q Exactive HF mass spectrometer (Figure 1A). Mixed species datasets have been used previously to validate protein quantification methods (Swift et al., 2013a), but the method has practical applications, such as when quantifying protein content in xenograft models (Ivanovska et al., 2017; Swift et al., 2013b). Mixed mouse/human samples provide a difficult problem for prospective quantification algorithms as approximately 70% of tryptic peptides are shared between mouse and humans and thus have to be discarded to obtain estimates for specific proteins. This meant that the quantification algorithms examined here had a limited number of observations with which to work. Secondly, the dynamic range of absolute protein abundances was derived from actual biological samples and so provided an accurate report of the error associated with estimating fold changes in low abundance proteins. Finally, this experiment yielded twin datasets (mouse and human); one with pronounced donor-donor variability (human cells) and one where all animals had the same genotype and environment (mouse skin), and so could be adequately explained by the model in equation (2) without donor effects. We reasoned that an algorithm that could account for donor-donor variability should be as accurate on the human dataset as on the mouse dataset.

**Figure 1.**
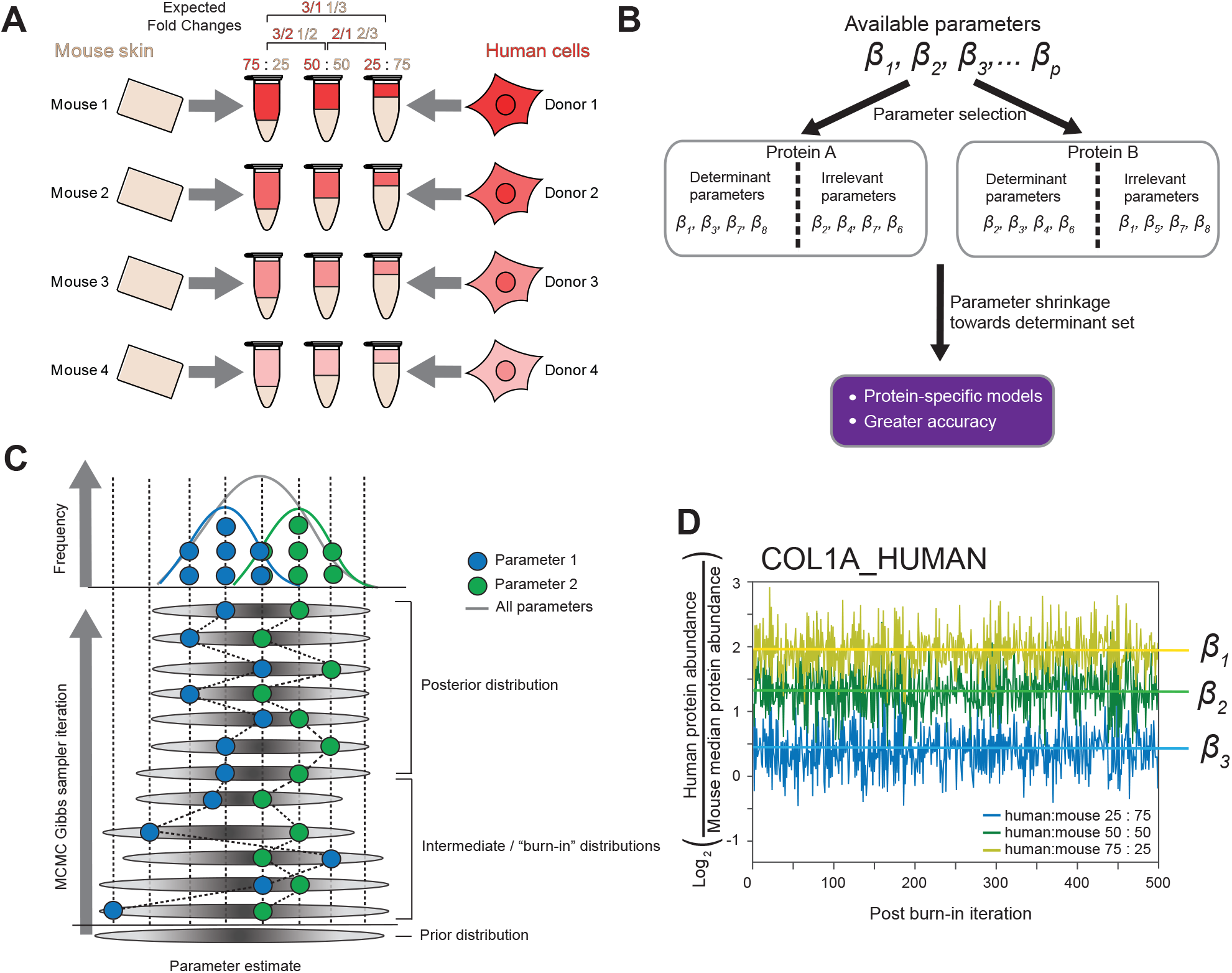
Extraction and preparation of mouse and human proteins for use in label-free quantification using multiple regression models. (**A**) Schematic diagram of the protocol used to prepare samples for the mixed species benchmark experiment. (**B**) Flow diagram showing the effect of regularisation on estimated parameters used in fitting models. (**C**) Schematic representation of how the Gibbs sampler arrives at the final (posterior) distribution for a single parameter. As the sampler proceeds, it will ideally converge on a single posterior distribution, as progressive iterations are biased towards areas of higher likelihood. Iterations from before this point are typically discarded (“burn-in” iterations) as those samples will not come from the target posterior distribution. Each parameter is then summarized from its respective estimated distribution (green and blue distributions) and all parameters lie on the target posterior distribution (grey distribution). (**D**) Example traces from post-burn-in samples to build posterior β distributions corresponding to log_2_ (protein abundance) estimates (denoted *β*_1_, *β*_2_ and *β*_3_). Traces show good mixing with no bias towards particular regions of their respective posterior distributions. Note that chains may not necessarily converge on exact expected fold change values, because: (i) the experimenter must first select one group to normalise the others to, so as to allow relative quantification; and (ii) imperfect accuracy.

To automatically select appropriate *β* parameters it was possible to employ regularisation strategies such as ridge (as implemented previously (Goeminne et al., 2016) and here as LME-H), LASSO or elastic net regression to select the best set of *β*s by shrinking unimportant *β*s to zero (Figure 1B). As LASSO has been shown to lack consistency when selecting from several correlated *β*s, we chose to test elastic net-based regularisation (implemented as BayesENproteomics) as the introduction of L_2_ regularisation has been shown to introduce consistency to LASSO estimates by inducing grouping of correlated *β*s (Zou and Hastie, 2005). However, the standard formula for calculating standard errors (SE) of *β* estimates (*σ*^2^ (***X***^T^***X***)^−1^, where *σ*^2^ is the residual variance and ***X*** is the design matrix, with superscript *T* denoting the transpose of ***X*** and SEs for each *β*, lie on the main diagonal of the resulting square matrix) was not suitable for sparse regression models due to shrinking of variance estimates by regularisation. This resulted in underestimation of variance and incorrectly lower SEs and p-values. While there were no simple ways to calculate SEs of coefficients in sparse regularised models, bootstrapping has commonly been used, however this is computationally intensive to do for hundreds of proteins. Bootstrapped SE estimates also differ from iteration to iteration, due to the inherent randomness of bootstrapping, thus compromising reproducibility. Bayesian regression methods addressed this limitation by estimating parameters as distributions for which summary statistics (e.g. means, standard deviations and confidence intervals) were trivial to calculate. Bayesian regression methods were also more flexible in how the model parameters and hyperparameters could be defined. This made Bayesian regression methods generally better suited to fit non-linear models, such as in equation (2), which frequentist methods may have struggled to fit. To implement this non-linear regression, we used a Monte Carlo Markov Chain (MCMC)-based Gibbs sampler to repeatedly sample from and update the prior conditional distributions for the parameters we wished to estimate (Figure 1C). Examples of the resulting chains are shown in Figure 1D and demonstrate good mixing and exploration of the posterior distributions.

A common assumption made is that the majority of missing values in bottom-up proteomics are assumed to be missing non-randomly (MNR) due to low abundance (Goeminne et al., 2016). To account for this lost variance, it is common to perform DGD imputation. In contrast, values can also be missing at random (MAR) and imputed from the observed values through local similarity-based approaches such as K-nearest neighbours.

The proportion of MNR to MAR values is strongly dataset- and even protein-dependent (Lazar et al., 2016), meaning that the optimal choice of distribution to impute from likely differs between datasets and proteins. Many state-of-the-art imputation methods rely on the user deciding whether they think that all missing values in a dataset are MAR or MNR and selecting an imputation method that performs well under these conditions (Lazar et al., 2016; Webb-Robertson et al., 2015). The nature of peptide missingness may vary between proteins, so it is unlikely that one method is suitable for all datasets and all proteins within them. To address this, we employed an adaptive multiple imputation (AMI) strategy, within the Gibbs sampler described in Figure 1, where a logistic regression determines if specific peptides or treatments positively correlate with missingness. Those missing values associated with parameters that showed higher than average, positive correlation with missingness were deemed to be MNR and imputed from a truncated Gaussian distribution, or MAR and imputed from a Gaussian distribution otherwise within the main Gibbs sampler (similar to the model-based imputation described in Karpievitch et al., 2009; Koopmans et al., 2014; Li et al., 2011).

To interrogate the accuracy of each imputation method we created a synthetic dataset consisting of a single protein with 1000 peptides measured across 2 treatment groups (G1 and G2) with 6 replicates each. Values for G1 and G2 were populated from normal distributions with means of 0 and −2, respectively, and variance = 1, such that the ground-truth (log-scale) fold change, *G*2 − *G*1 = −2. Zero, 10, 20 or 30% of these values were deleted randomly (simulating MAR missingness) along with all values < −∞, −2, −1 or 0 (simulating MNR missingness) and we asked which imputation method could most accurately recapitulate the deleted values by comparing the root mean squared error (RMSE) between imputed values and ground truth deleted values. While no single method was capable of fully approximating the deleted values, DGD imputation showed the worst performance when MAR dominated and improved as MNR values became more prevalent (Figure 2A). As expected, KNN and BPCA showed the opposite trend with high accuracy in MAR-dominated datasets (especially for BPCA); decreasing as MNR proportion increased (Figures 2B, C). Model-based imputation using DanteR (Karpievitch et al., 2009) showed good performance with small proportions of missing values, but performed markedly decreased as missingness proportion increased (Figure 2D). AMI showed good accuracy in MAR-dominated datasets (typically intermediate between KNN and BPCA, depending on missingness proportion), but importantly, AMI was not negatively affected by the presence of MNR values and exceeded the performance of BPCA and KNN in cases of pure MNR missingness (Figure 2E). These results showed that AMI, as implemented in BayesENproteomics, possessed more consistent accuracy, and may be better suited to imputing values in a mixed-missingness-type dataset. Figure 3A shows estimation of log_2_ fold changes for non-differentially abundant proteins (339 with >2 unique peptides) in pairwise comparisons between three technical replicates from the same (mouse skin) sample, comparing the BayesENproteomics algorithm with other commonly used peptide-based models, namely: ordinary least-squares (OLS), as utilised in MSStats (Choi et al., 2014) with donor effects added (this manuscript) and ridge regression/mixed-effects models with Huber residual weights, (LME-H) as utilized in MSqRob (Goeminne et al., 2018, 2016) but necessarily modified to quantify donor effects, and peptide:treatment and peptide:donor interaction effects (see Methods section). All methods correctly determined log_2_ fold change values clustered around zero. Both LME-H and BayesENproteomics showed a single statistically significant false positive (p-value of false discovery, with Benjamini-Hochberg correction, BHFDR < 0.05) that disappeared when the imputation method was switched from DGD to AMI.

**Figure 2.**
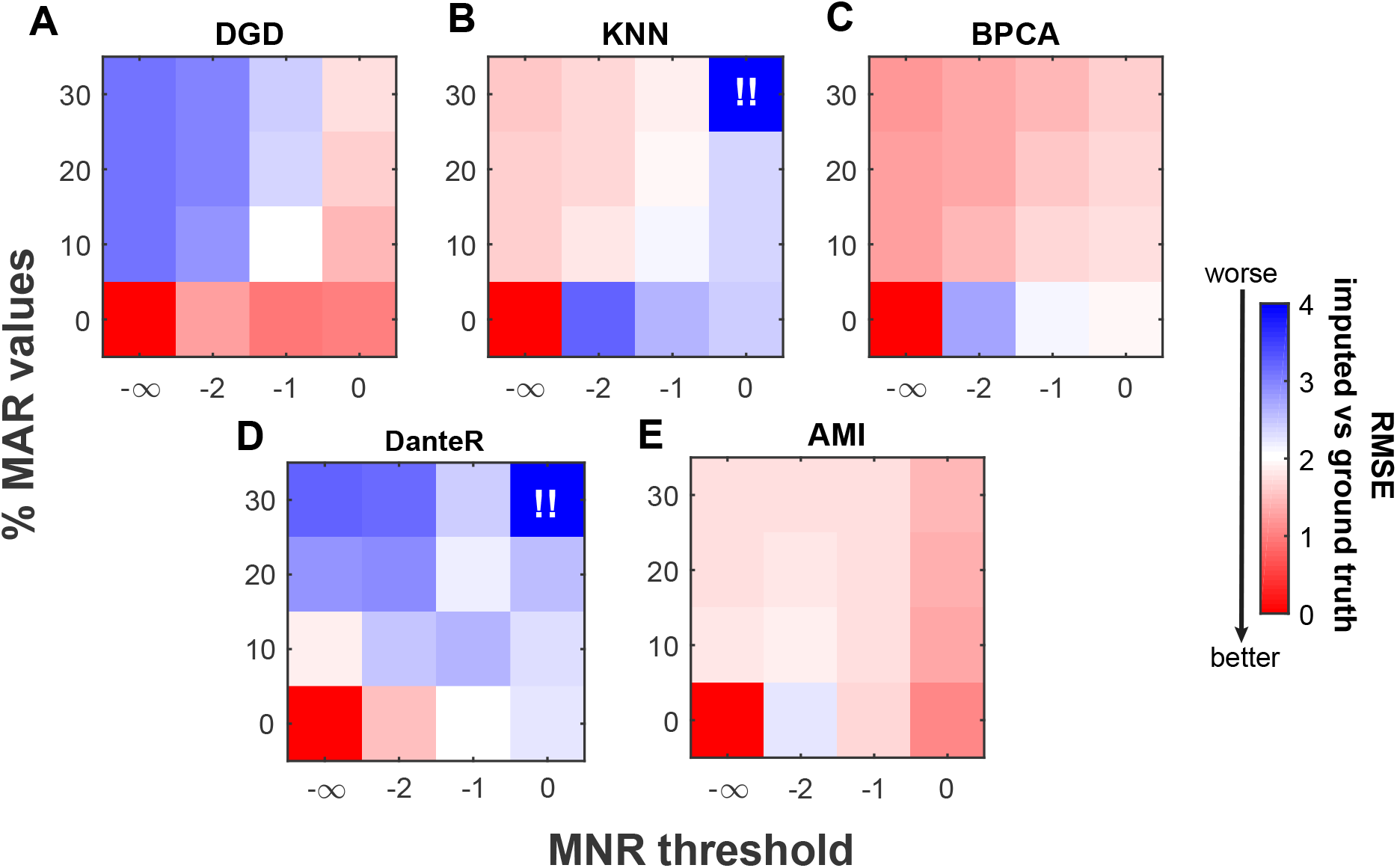
Comparison of missing value imputation strategies. Heatmaps showing RMSE of imputed values against ground truth deleted values for (**A**) DGD; (**B**) KNN (where K = 11); (**C**) BPCA; (**D**) model-based imputation using DanteR; and, (**E**) AMI. Colours closer to red denote smaller RMSEs and overall better performance. Synthetic dataset had 12000 values in total. To simulate MNR missingness values below the given threshold were deleted, corresponding to 0% (threshold = −∞), 1.2 % (−2), 8.9 % (−1) or 29 % (0) missing values. !! = failure of KNN and DanteR to run in this scenario due to too many missing values.

**Figure 3.**
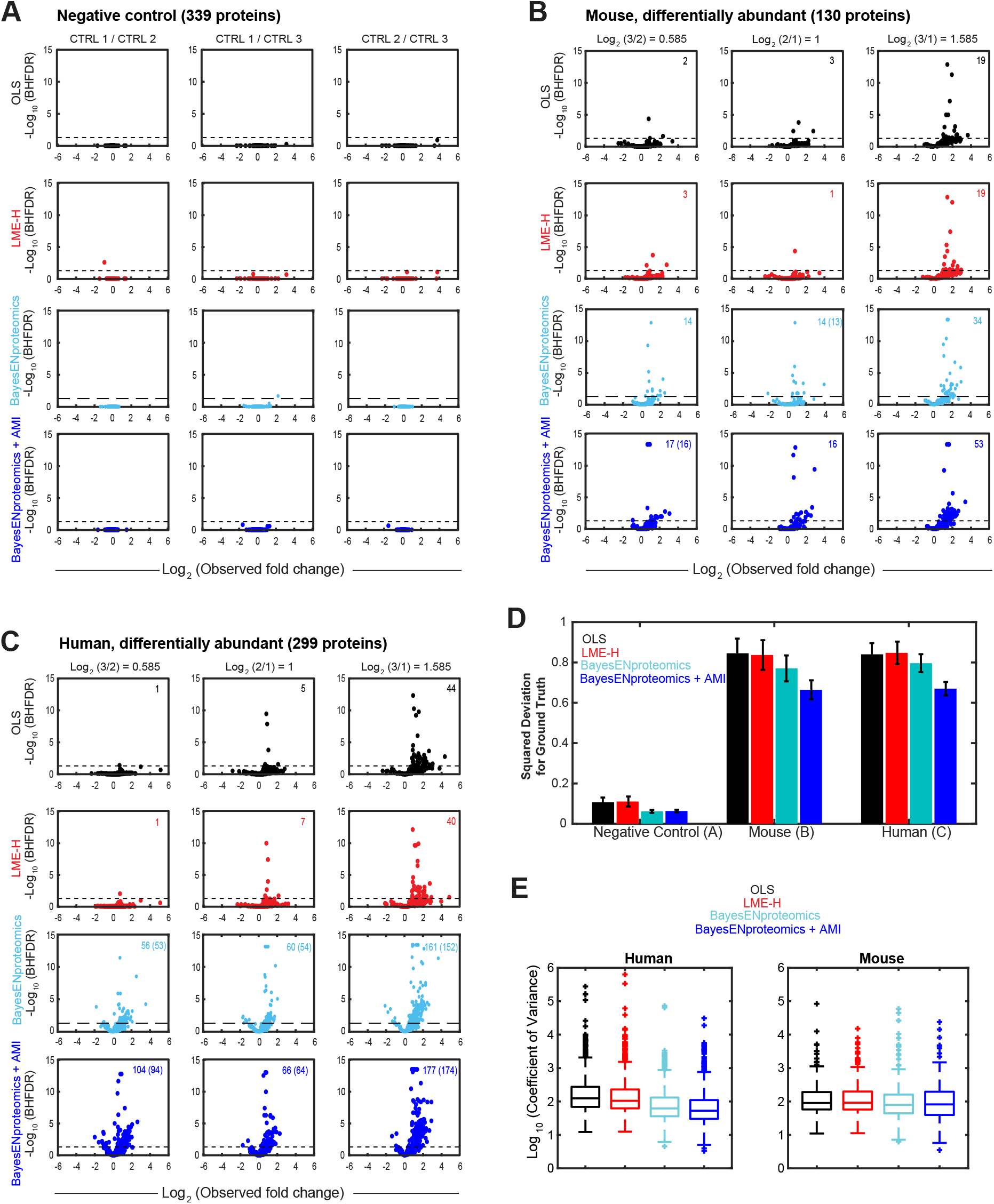
BayesENproteomics increases specificity compared to other multi-variate regression methods. (**A**) Volcano plots showing quantification of fold change estimates (x-axis) vs. significance (y-axis) observed in pairwise comparisons between three technical replicates. (**B-C**) Volcano plots showing quantification of mouse and human proteins, respectively, across all comparisons in the mixed human:mouse benchmark experiment. Numbers in top right corners denote numbers of true positives observed with numbers in brackets denoting true positives showing the expected direction of fold change. (**D**) Bar chart showing squared deviation from ground truth across all contrasts for the indicated dataset. Bars represent means ± standard error of squared deviations. (**E**) Boxplots showing coefficients of variation (CVs) for all differentially abundant proteins in the mixed human:mouse benchmark experiment. Dotted lines in (A)-(C) indicate Benjamini-Hochberg false discovery rate corrected p-value (BHFDR) = 0.05. OLS fit using simple model in equation (1), black; LME-H according to equation (2), red; BayesENproteomics as in (2), cyan; BayesENproteomics with AMI, blue. n = 4 human donors, 4 mice.

Estimation of differentially abundant mouse (130 proteins with >2 unique peptides; Figure 3B) and human (299 proteins with >2 unique peptides; Figure 3C) proteins in our mixed human:mouse dataset showed larger variation in observed fold change values, with a number of proteins giving consistently incorrect fold change directions, regardless of the regression method (Figure S1) – a common problem in label-free quantification (Zhang et al., 2017), even with state-of-the-art data-independent acquisition methods (Navarro et al., 2016). While BayesENproteomics improved specificity, it typically did not improve the accuracy of these consistently incorrect fold change estimates. Imputation method appeared to more strongly determine these incorrect fold change estimates, suggesting that incorrect estimates were due to missing values. Application of AMI corrected some incorrect fold change estimates but worsened others compared to DGD imputation, suggesting that pure MNR imputation may be more appropriate for a subset of proteins (Figure S1). OLS and LME-H models had difficulty detecting any significantly differentially abundant proteins for expected fold changes < 3 (Figures 3B, C). In contrast, BayesENproteomics correctly identified more proteins as significantly differentially abundant in all comparisons (Figures 3B, C), with AMI increasing the number of true positives over DGD imputation. Importantly, fold change estimates calculated by BayesENproteomics possessed lower mean squared deviation from ground truth compared to those from OLS and LME-H, which improved further with AMI (Figure 3D). BayesENproteomics also showed decreased coefficient of variance (CV) values for differentially abundant proteins in the mixed human:mouse dataset compared to both OLS and LME-H (Figure 3E).

### BayesENproteomics correctly detects increases in stable isotope-labelled proteoforms following PNGase F treatment in H_2_^18^O

Protein function and activity is strongly determined by diverse PTMs, giving rise to different proteoforms within a given protein population. MS-based proteomics represents a powerful means to systemically interrogate changes in PTM abundance. However, determining relative PTM fold changes from PTM’d peptide abundance is dependent on accurate protein quantification (Wu et al., 2011). While it is possible to calculate protein abundance from only unmodified peptides, this ignores the fact that some proteins are constitutively modified as part of their normal maturation pathway (e.g. many extracellular matrix proteins are heavily glycosylated prior to secretion) and excluding these PTM-containing peptides from protein quantification would exclude a large proportion of all proteoforms present for that protein. While methods have been developed to analyse individual proteoform abundance in multiplexed, labelled experiments (Malioutov et al., 2017), labelling is not always feasible and methods for quantifying differentially abundant PTMs in label-free experiments lag behind. Furthermore, while an ideal scenario would be to directly compare modified to unmodified peptide ratios (Tsai et al., 2017), this is not always achievable in label-free proteomics of complex samples. While linear regression modelling has been shown to impart greater power and accuracy to protein quantification compared to summary statistical methods (Clough et al., 2009; Goeminne et al., 2015), applying it to study PTMs is difficult due to the requirement for complex non-linear models that may be subject to overfitting.

To interrogate how well the three models implemented here were able to detect differentially abundant PTMs, we developed a benchmark dataset using PNGase F to introduce a stable isotope (^18^O) label on N-linked glycosylated residues (Figure 4A). We reasoned that PNGase F-treated peptides should show an increase in N-linked ^18^O labelling compared to spontaneous deamidation observed in control peptides incubated in H_2_^18^O buffer alone. To test if our PNGase F treatment of peptides had worked correctly (independently of any subsequent quantification methods) we first searched treated and control samples separately and counted the total number of N- and Q-linked, ^18^O-labelled peptides identified (Figure 4B). This analysis showed that more ^18^O-labelled peptides were identified in the PNGase F-treated samples than controls, which contained 79 N-linked peptides presumably from spontaneous non-enzymatic deamidation. We expected that most spontaneously deamidated peptides would be unaffected by PNGase F treatment and that we would only observe an increase in relative ^18^O-peptide abundance or parent ^18^O-protein abundance for PNGase F-responsive peptides. Furthermore, as spontaneous deamidation can occur to both Q and N residues, resulting in ^18^O-E and ^18^O-D residues, respectively, while PNGase F catalyses only deamidation of N-linked glycosylation sites, comparison to fold changes of Q-linked deamidation provided an ideal negative control.

**Figure 4.**
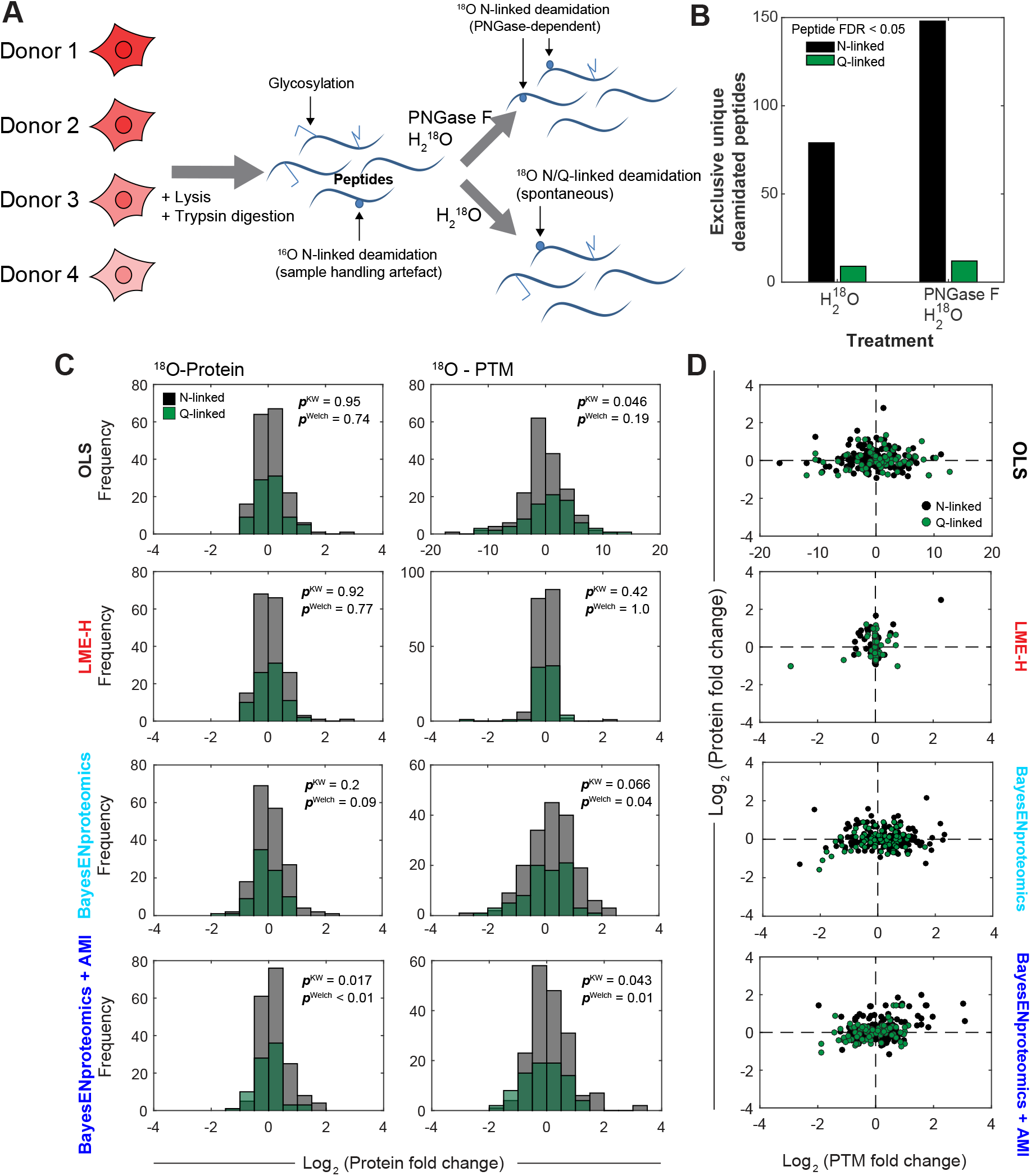
PNGase-catalysed stable isotope labelling quantified by summarization with normalisation and multivariate regression methods. (**A**) Schematic diagram illustrating experimental setup for the variable PTM benchmark dataset. (**B**) The number of deamidated peptides was increased following 5 hours PNGase F treatment. Only peptides from proteins identified by >1 peptide were counted. (**C**) Histograms showing fold changes in N-linked ^18^O labelled N-linked deamidation (yellow) and ^18^O labelled Q-linked deamidation (green). (**D**) Scatter plots comparing protein and PTM log2 fold changes for the indicated quantification methods. n = 5 human donors. Indicated p-values show significance from comparison of N-linked (yellow) and Q-linked (green) histograms calculated by Welch’s two-sample t-test (p^Welch^) or Kruskal-Wallis test (p^KW^).

To detect differentially abundant PTMs we employed three statistical methods. The first was a simple summary statistical extension to the OLS protein quantification method wherein log_2_ fold changes for donor-normalised PTM’d peptide intensities were normalised to the log_2_ fold changes of their parent protein (calculated by OLS). The second involved using the ridge regression performed by LME-H to fit the non-linear model described in equation (2) where peptide:group and peptide:donor interaction *β*s were entered as random intercept terms to limit overfitting. Finally, we used BayesENproteomics to fit the non-linear model described in equation (2) to calculate PTM fold changes.

From simple cell lysates, following PNGase F treatment without enrichment we found 261 ^18^O-labelled sites (180 N-linked, 81 Q-linked) in 172 proteins when samples from all groups were aligned and search together. Histograms in Figure 4C showed that OLS and LME-H could not discern an increase in N-linked ^18^O-protein abundance following PNGase F treatment compared to Q-linked (all p-values > 0.05). In contrast, BayesENproteomics was able to correctly discern a positive skewing of the N-linked ^18^O-protein histogram compared to that of the Q-linked proteins that improved to significance with concomitant application of AMI (all p-values < 0.05, Figure 4C).

Similarly, when measuring differentially abundant PTMs there was great disparity between the three methods. The summary statistical extension to OLS produced noisy estimates of PTM fold changes and was unable to discern an increase in N-linked ^18^O-PTMs following PNGase F treatment compared to Q-linked (Figures 4C, D). LME-H displayed overly strong regularisation of interaction terms, resulting in the majority of PTM fold change estimates being approximately zero (Figures 4C, D). BayesENproteomics estimates of N-linked ^18^O-PTM fold changes were correctly positively skewed (Figures 4C, D) and significantly different from Q-linked ^18^O sites with AMI. This showed that BayesENproteomics with AMI could quantify PTM fold changes more accurately than either normalised summary statistics or mixed-effects models.

### Pathway analysis using linear modelling (PALM)

Theoretically, modelling pathway activity as a function of protein abundances (PALM) can have several advantages over classical pathway over-representation (Figure 5A) and enrichment (Figure 5B) analysis. Similar to enrichment analysis, the PALM approach proposed here (Figure 5C) utilises all proteins regardless of significance or fold change magnitude. This avoids potentially arbitrary decisions used in defining “interesting” and “null/background” sets for over-representation analysis (reviewed in Huang et al., 2009) that can become difficult if a dataset is further sub-divided (e.g. experiments where proteins are divided into several clusters according to changing expression profiles). Secondly, unlike enrichment analysis, PALM does not assume that there is no intrinsic correlation between proteins in a given dataset. This assumption can be violated in pull-down or proximity-labelling experiments where a specific subset of proteins is quantified. Additionally, PALM uses additional information regarding the direction and magnitude of fold changes and the error regarding their estimation to provide an overall estimate of how a given pathway behaved based on information available in a given dataset (Figure 5D). Finally, PALM provides pathway-level fold change and error estimates, facilitating pathway-level clustering in complex, multi-treatment or time-series datasets to gain a better “bird’s-eye view” of what is happening in a dataset. We compared Reactome (Fabregat et al., 2018; Milacic et al., 2012) pathway-level fold changes estimated using either the Reactome website tool (https://Reactome.org) or PALM. Pathway-level fold changes were estimated from the technical replicate and mixed human:mouse datasets using BayesENproteomics protein-level fold change estimates (Figures 3A-C). PALM pathway fold change estimates correlated with pathway averages calculated by Reactome (Figure 5E). While the magnitude of pathway-level log_2_ fold changes was typically not equal to their protein components due to being inferred from relatively fewer observations and down-weighted according to uncertainty in initial protein fold change estimates (always > 0), directionality was conserved leading to an estimate of whether that pathway significantly increased or decreased. PALM BHFDR-adjusted significance estimates of differentially abundant pathways showed an FDR of 0.03 (4 false positives out of 120 non-significant comparisons); this was acceptable within the pre-specified BHFDR cut-off of 0.05 (Figure 5E). Enrichment analysis using PANTHER (pantherdb.org; Mi et al., 2017) showed no significant enrichments for any of the comparisons in Figure 5E (Supplementary Tables S1-3), likely because of the high between-protein correlation within the mouse:human dataset where most proteins had similar fold changes, resulting in a “null” set with negligible difference to any given “interesting” set.

**Figure 5.**
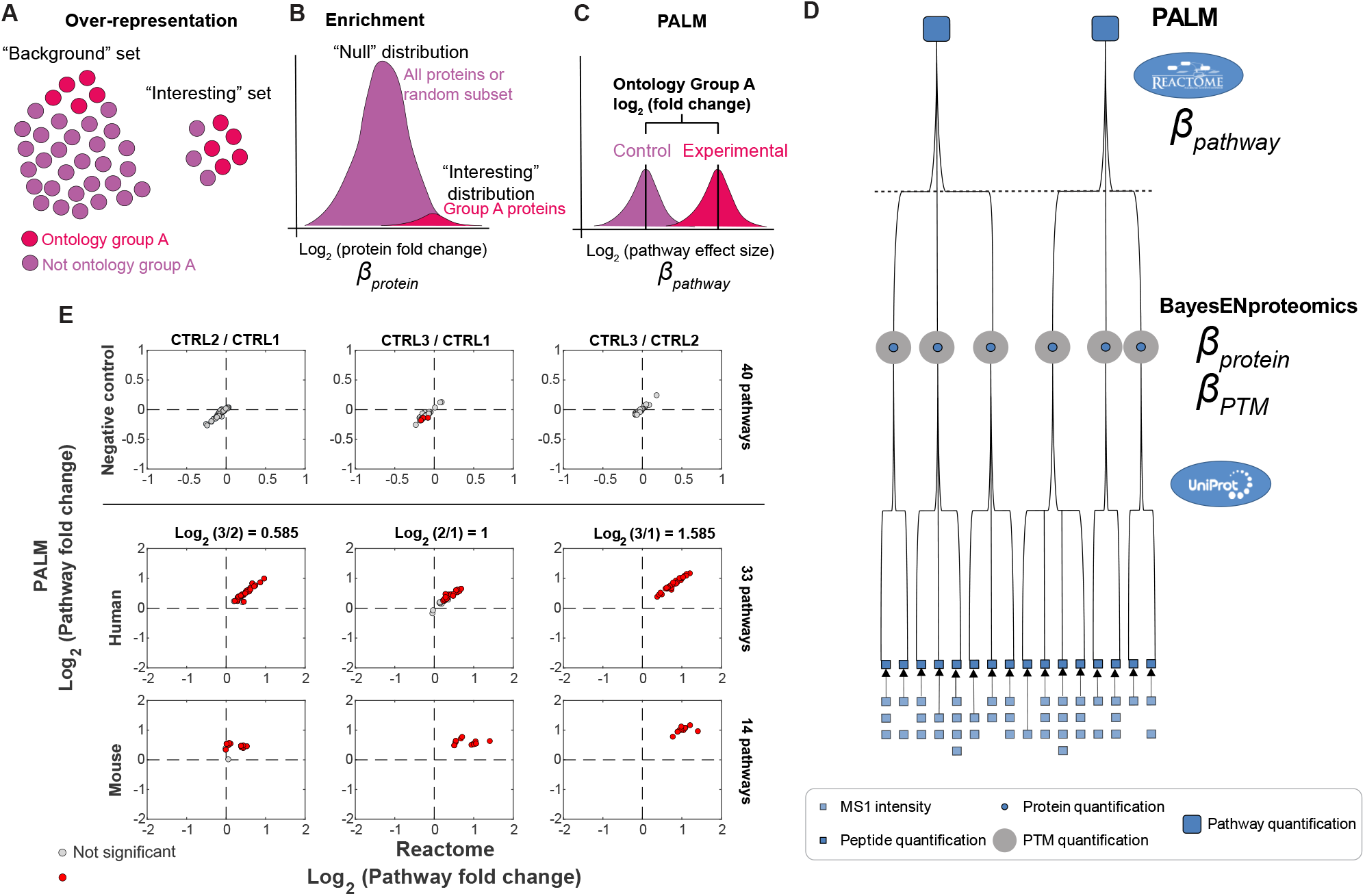
Pathway analysis using linear models (PALM) on mixed-species dataset. (**A-C**) Schematic diagrams representing different methods of pathway analysis. The advantages of PALM over classical enrichment and over-representation analysis are: (i) no need to select “interesting” and “null/background” sets; (ii) incorporation of uncertainty of protein fold change estimates into pathway-level fold changes. (**D**) Schematic diagram showing hierarchical tree structure for linear model assembly from MS1 intensities (bottom) to pathways (top). *β*s correspond to effect sizes estimated in equations 1, 2 and 16. (**E**) Scatter plots showing pathway-level fold changes calculated from the BayesENproteomics with AMI protein-level fold changes for the technical replicate negative control (top) and mouse:human mixed species datasets (middle, bottom) from Figures 3A-C, using either Reactome’s own tool (x-axis) or PALM (y-axis). As Reactome.org only provides significance for over-representation given a set of protein identifiers, a statistical power comparison is not possible. Enrichment analysis using PANTHER showed no significant differences for any pathways. Each dot represents a single Reactome pathway. Pathways follow the expected direction of the component proteins, though confidence in original protein-level fold change estimates may affect exact magnitudes of pathway-level fold changes. Only pathways with ≥5 proteins present in the dataset are shown.

## Discussion

Multivariate regression methods represent a powerful means to deconvolute sources of variance in complex experiments. Here we attempted to design a statistical method that was able to accurately quantify protein and PTM fold changes in label-free proteomics experiments performed on heterogeneous primary human samples. Others have previously shown the superiority of linear regression-based effect size estimates over summary statistics (Clough et al., 2009; Goeminne et al., 2015). Here we outlined a weighted Bayesian regression algorithm with elastic net regularisation to cope with differentially abundant PTMs as well as the high variability and generally low availability of samples from human donors, while still maintaining high interpretability of the resulting models (Figure 6). In particular, BayesENproteomics incorporates an “adaptive” missingness mechanism that attempts to determine whether a value is MAR or MNR. Missing value imputation is performed as part of the Gibbs sampler and is thus unfortunately not available as a standalone component. In addition, we utilised a novel weighing method that considered confidence in peptide identification to improve quantification accuracy. We demonstrated the accuracy and specificity of the algorithm using serial mixed species datasets and stable isotope labelling using peptides obtained from multiple human donors.

**Figure 6.**
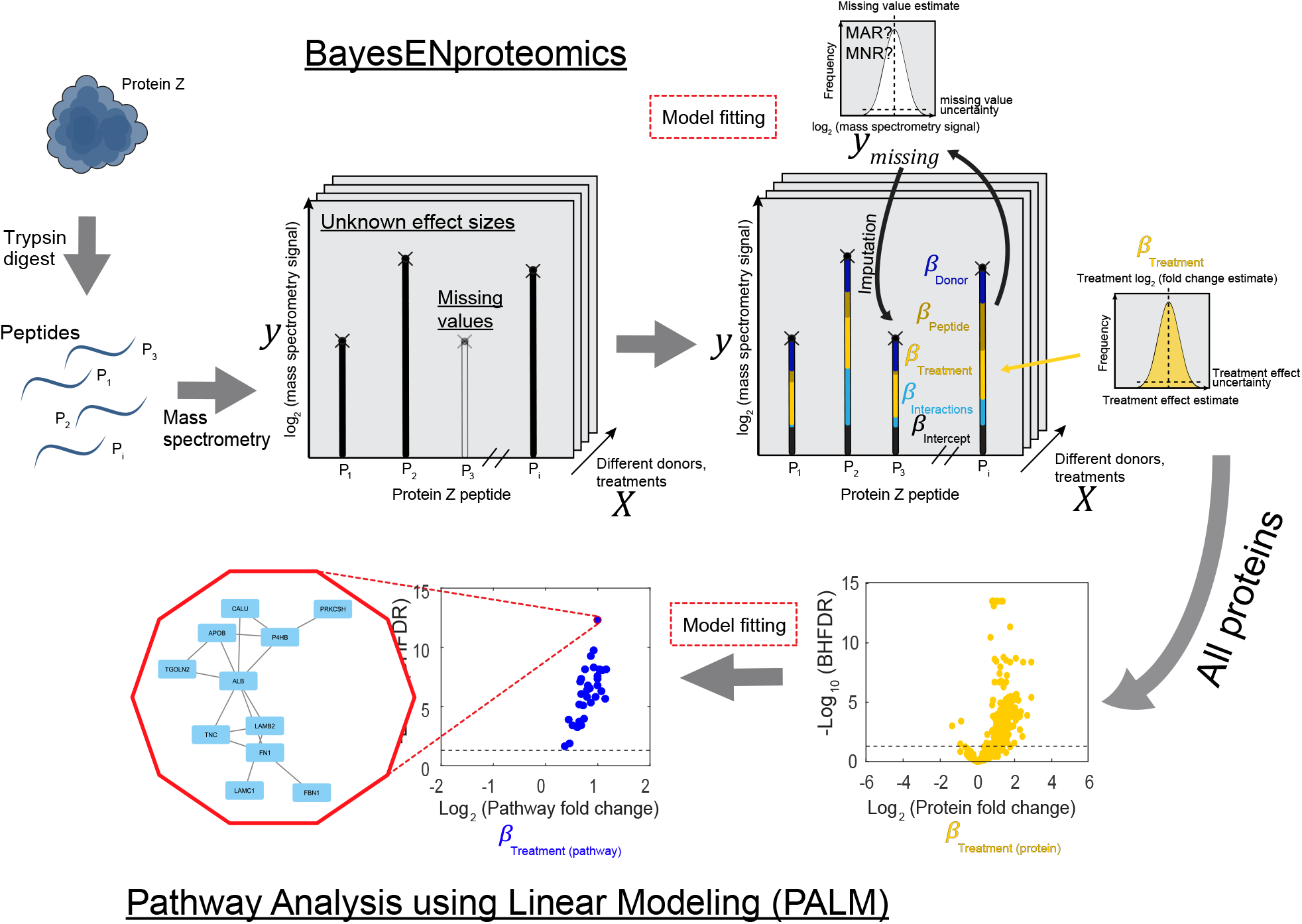
Summary schematic of complete BayesENproteomics and PALM workflow, including missing value imputation using AMI. Symbols correspond to those in equations 1 and 2. Regression modelling aims to decompose experimental variability contained in logged signal intensity (*y*, with *X* denoting which observation in *y* came from which treatment, peptide, donor, etc) into constituent parts (*β*s), some of which may be of interest to the experimenter (e.g. variability caused by experimental treatment, *β*_*Treatment*_) and others that – while less interesting – may still need to be accounted for (e.g. variability caused by peptide behaviour during mass spectrometry, *β*_*Peptide*_). BayesENproteomics achieves this using a Gibbs sampler to iteratively optimize *β* estimates, resulting in posterior distributions for both *β*s and any missing values (*y*_*missing*_) in the dataset. Protein-level logged fold changes (*β*_*Treatment (protein)*_) are then fed into a second round of model fitting to provide pathway-level logged fold changes (*β*_*Treatment (patway)*_).

Characterisation of complex samples is notoriously complex due to ionisation suppression and detection masking by peptides with similar m/z ratios often leading to missing values and underestimation of larger fold changes (this study and Goeminne et al., 2016). The mixed species dataset used here represented an especially difficult case with most (~70%) peptides rendered unusable due to being shared between the two species. While all methods tended to underestimate fold change magnitude (likely due to conflicting peptides in data-dependent analysis, exacerbated in mixed species analysis), we showed that BayesENproteomics was better able to confidently identify differentially abundant proteins and PTMs than simpler regression methods with greater accuracy in fold change estimates compared to OLS and LME-H regression models. Furthermore, we show that missing value imputation was a strong determinant for those proteins with consistently incorrect fold change estimates (poor accuracy). BayesENproteomics typically did not improve the accuracy of fold change estimates for these proteins, but was still able to classify them as significantly different in agreement with ground truth. This increased specificity without increased accuracy could potentially be dangerous, highlighting the importance of subsequent experimental validation and may warrant exclusion of proteins with proportions of missing values above some pre-set threshold.

We used the iterative nature of MCMC processes to incorporate multiple imputation of missing values into protein fold change quantification. This eliminated the need for separate imputation and quantification steps and allowed final fold change error estimates to consider the intrinsic uncertainty regarding imputation of missing values (Schafer and Olsen, 1998). Here we employed a standard random imputation from prior distributions, but this could easily be substituted for any desired method.

Currently, BayesENproteomics does not utilise shared peptides, an approach that others have shown improves accuracy (Blein-Nicolas et al., 2012) at the cost of increased computation time. Use of empirical Bayes correction of variances (as in all methods tested here) can increase power by borrowing information from all proteins/PTMs without having to fit a single model for all proteins simultaneously (as would be required to allow the use of shared peptides) for the purposes of calculating significance. MCMC samplers are known to be particularly computationally intensive, particularly if the experimenter wanted to extend it with additional fixed interaction effects to quantify PTM behaviour. BayesENproteomics side-steps this problem by fitting individual models for each protein, at the cost of not being able to use shared peptides. On an especially rich dataset, such as those generated by a modern Q Exactive HF mass spectrometer, analysis using BayesENproteomics can take several hours to process on a standard desktop computer. This is approximately three times longer than LME-H and several times longer than OLS regression. Importantly, the use of Gaussian/Laplacian priors for estimating *β* parameters presumed that logged fold changes lay on an approximately normal distribution. If it was suspected that this was not the case, BayesENproteomics may not be a suitable method of analysis.

Linear models are extremely customisable, and indeed, can require significant tailoring to best analyse complex multivariate experiments. Without sufficient background knowledge it can be very difficult to anticipate all potential sources of variability that need to be accounted for in any putative statistical model. Nevertheless, BayesENproteomics showed negligible perturbation by inclusion of potentially extraneous parameters and so is well-suited for complex experimental setups that may include different batches, fractions, serial extractions, instruments and donors. Regularisation hyperparameters for additional effects can be sampled from the conditional distributions detailed here.

Finally, we extended our linear modelling pipeline to encompass pathway analysis. In transcriptomics experiments it is common to achieve coverage of the entire transcriptome (> 20000 transcripts in a given human cell/tissue), where hundreds to thousands may be differentially regulated in response to any given treatment. These large numbers make over-representation analysis using χ^2^ or Fisher’s exact tests well-suited to analysing differentially regulated transcriptomic processes. In contrast, proteomics experiments typically cover only a fraction of the total proteome (< 3000 proteins out of a potential > 20000 in a given human cell/tissue, increased further when considering multiple proteoforms) where only a hundred or fewer proteins may be differentially regulated, making it difficult to discern statistically significant process enrichment and erroneously giving the impression that “nothing is happening”, often despite obvious morphological or metabolic changes assessed by other methods. Attempting to model pathway-level changes as a function of component parts for transcriptomics has previously shown that incorporating fold change estimates can increase specificity for discriminating pathway-level changes (Ozerov et al., 2016). Although, it should be noted that the aim with PALM is not biomarker discovery (for which more stringent statistical cut-offs are needed) but rather to simply distil a complex dataset with many proteins down to fewer, more readily understandable pathways by assessing what processes are represented and what they do in response to a given treatment. This works in a similar way to previous enrichment tools (e.g. PANTHER (Mi et al., 2017)) and is comparable to previous implementations of Bayesian regression in proteomics (Henao et al., 2013), however here we opted to maintain interpretability by utilizing curated protein-pathway annotations when picking proteins to use in pathway-level model fitting. Here we chose to use Reactome pathway annotations (Fabregat et al., 2018), but alternative categorical annotations could be used if desired. PALM provides a way of compressing information from many protein/proteoform fold changes into fewer annotated pathway fold changes, aiding interpretation of complex, multi-dimensional datasets. PALM can be used as a standalone pathway analysis tool; although it does assume protein fold changes are formatted according to the BayesENproteomics protein-level output format (see examples on https://github.com/VenkMallikarjun).

With the improvement of MS technology, and the production of highly efficient unconstrained searching algorithms (Devabhaktuni et al., 2018; Kong et al., 2017), identification and quantification of increasingly rare PTM’d peptides, and thus their parent proteoforms, has become possible even without enrichment. Many quantification methods have discarded or otherwise diminished the influence of these potentially relevant peptides in bottom-up protein quantification. Here, the use of regularised interaction coefficients facilitated the reporting of differentially abundant PTMs and the exclusion of potentially misidentified peptides from protein (and subsequent pathway) fold change estimates. Global quantification of proteoforms represents an exciting avenue for understanding how cell function is regulated post-translationally. BayesENproteomics was able to account for differences in PTM’d peptide behaviour whilst increasing accuracy of protein quantification in benchmark datasets with limited observations and inter-donor variability.

## Materials and Methods

**Table 1.** Reagents and Tools Table.

| Reagent/Resource | Reference or Source | Identifier or Catalogue Number |
| --- | --- | --- |
| **Experimental Models** |  |  |
| C57BL/6J (*M. musculus*) skin | Skin provided by Hardman group, mice originally from Jackson Lab | 000664 |
| Primary human mesenchymal stromal cells (isolated from patient bone marrow) | Wrightington Hospital, Wigan, UK | N/A |
| **Chemicals, Enzymes and other reagents** |  |  |
| PNGase F from *Elizabethkingia meningoseptica* | Sigma-Aldrich | P7367 |
| H<sub>2</sub><sup>18</sup>O | Sigma | 329878 |
| low-glucose DMEM with pyruvate | Thermo Fisher Scientific | 11885084 |
| fetal bovine serum | Labtech.com |  |
| penicillin/streptomycin cocktail | Sigma-Aldrich | P4333 |
| ammonium bicarbonate | Sigma-Aldrich | 09830 |
| sodium dodecyl sulphate | Sigma | L3771 |
| sodium deoxycholate | Sigma | D6750 |
| protease inhibitor cocktail | Sigma | P8340 |
| phosphatase inhibitor cocktail | Sigma | P0044 |
| dithiothreitol | Sigma | D0632 |
| immobilized trypsin beads | Perfinity (via Thermo Fisher Scientific) | 60109-101 |
| urea | Thermo Fisher Scientific | 29700 |
| calcium chloride dihydrate | Sigma | C3306 |
| iodoacetamide | Sigma | I1149 |
| trifluoroacetic acid | Sigma | 61030 |
| ethyl acetate | Sigma | 34858 |
| POROS R3 beads | Thermo Fisher Scientific | 1133903 |
| acetonitrile | Thermo Fisher Scientific | TS-51101 |
| formic acid | Thermo Fisher Scientific | 28905 |
| **Software** |  |  |
| Matlab (2012a or higher) with Bioinformatics and Statistics and Machine Learning toolboxes | The MathWorks Inc. |  |
| Bayesian PCA imputation code | (Oba et al., 2003) | http://ishiilab.jp/member/oba/tools/BPCAFill.html#i7 |
| BayesENproteomics (Matlab code) | This manuscript | https://github.com/VenkMallikarjun/BayesENproteomics |
| Progenesis QI for proteomics (v3.0) | Non-linear Dynamics |  |
| Mascot Daemon | Matrix Science | http://www.matrixscience.com/ |
| Scaffold (v3.0) | Proteome Software, Inc. | http://www.proteomesoftware.com/products/scaffold/ |
| R 3.5.2; RStudio 1.1.534 | R Consortium | https://www.rstudio.com/ |

### Methods and Protocols

#### Cell culture

For the mixed species benchmark experiment, primary human mesenchymal stem cells (MSCs) were isolated from the bone marrow (knee and hip) of a single donor with informed consent and ethical approval, using established protocols (Strassburg et al., 2010). All experiments were performed in accordance with relevant guidelines and regulations, and with National Research Ethics Service and University of Manchester approvals. MSCs were cultured on tissue culture treated polystyrene (TCTP) in low-glucose DMEM with pyruvate (Thermo Fisher Scientific) supplemented with 10% fetal bovine serum (FBS, Labtech.com) and 1% penicillin/streptomycin cocktail (PS, Sigma-Aldrich).

#### Preparation of primary human MSCs lysates

Cells were lysed in 25 mM ammonium bicarbonate (AB) buffer containing 1.1% sodium dodecyl sulphate (SDS, Sigma), 0.3% sodium deoxycholate (Sigma), protease inhibitor cocktail (Sigma), phosphatase inhibitor cocktails (Sigma). Six 1.6 mm steel beads (Next Advance) were added to the tube and samples were homogenised with a Bullet Blender (Next Advance) at maximum speed for 2 minutes. Homogenates were cleared by centrifugation (12 °C, 10000 rpm, 5 minutes).

#### Preparation of mouse skin lysates

Skin from 3-month old female C57BL/6J mice was a gift from the Hardman group (University of Manchester; all animal work was performed under UK Home Office and local ethics committee approval). Ventral skin was shaved and scraped, then dissected into 1 mm^3^ pieces. 100 *µ*L of 8 M urea (Fisher Scientific) in 25 mM AB containing 25 mM dithiothretol (DTT, Sigma), protease and phosphatase inhibitor cocktails (Sigma) was added to the tissue sections with six 1.6 mm steel beads (Next Advance). Samples were then homogenised with a Bullet Blender (Next Advance) at maximum speed for 3 minutes. Resulting homogenates were cleared by centrifugation (12 °C, 10000 rpm, 5 minutes).

#### Preparation of tryptic peptides

Immobilised trypsin beads (Perfinity Biosciences) were suspended in 150 *µ*L of digest buffer (1.33 mM CaCl_2_, Sigma, in 25 mM AB) with 50 *µ*L of cell or tissue lysate and shaken at 1400 rpm overnight at 37 °C in a thermocycler. The resulting digest was then reduced by the addition of 4 *µ*L of 500 mM DTT (Sigma, in 25 mM AB; 10 min. shaking at 1400 rpm at 60 °C) and alkylated by the addition of 12 *µ*L 500 mM iodoacetamide (Sigma, in 25 mM AB; 30 min. shaking in the dark at room temperature). Immobilised trypsin beads were removed by centrifugation at 10000 rpm for 10 min. Supernatant containing reduced, alkylated peptides were transferred to 1.5 mL ‘LoBind’ Eppendorf tubes and acidified by addition of 5 *µ*L 10% trifluoroacetic acid (TFA, Sigma) in water, and cleaned by two-phase extraction (3 × addition of 200 *µ*L ethyl acetate, Sigma, followed by vortexing and aspiration of the organic layer). Peptides were desalted using POROS R3 beads (Thermo Fisher). Briefly, R3 beads were thoroughly mixed in 50 % (v/v) acetonitrile (Sigma; prepared in milliQ H_2_O; elute solution) by vortexing, to a concentration of 10 mg/mL. For each sample, 120 *µ*L of bead suspension was pipetted into a well of a 96-well filtration membrane plate. The plate was then centrifuged (200 × g, 1 min, room temperature) and the run-through was collected into a separate 96-well plate. The beads were then resuspended in 50 *µ*L elute solution and centrifuged again twice before repeating the procedure with 0.1 % TFA (prepared in milliQ H_2_O; wash solution). The run-through from the collection plate was discarded and the R3 beads were resuspended in 100 *µ*l of sample before centrifugation. This process was repeated until all sample had been added. Bead-bound peptides were washed twice in wash solution. The run-through was discarded and the collection plate was washed with elute solution. Peptides were then eluted by two rounds of suspension in 50 *µ*L elute solution and centrifugation. Eluted peptides were then lyophilized. Peptide concentrations (measured by Direct Detect spectrophotometer, Millipore) in injection solution (5% HPLC grade acetonitrile, Fisher Scientific, 0.1% TFA in deionized water) were adjusted to 200 ng/*µ*L prior to MS analysis.

#### Preparation of serial mixtures of human and mouse peptides

Primary human MSCs and mouse skin sections were prepared separately as described above. Tryptic peptides from each preparation were mixed in ratios of 3:1, 1:1 and 1:3 (human:mouse, one series for each human donor:mouse), as described previously in (Swift et al., 2013a).

#### Stable isotope labelling of peptides

Desalted peptides were resuspended in 100 mM AB dissolved in H_2_^18^O with or without 5 U PNGase F (Sigma), to a final volume of 10 *µ*L. Peptides were then incubated in a thermocycler (1400 rpm, 37 °C) for 5 hours before drying down to a minimal volume in a vacuum centrifuge and resuspension in injection solution prior to MS analysis.

#### Mass spectrometry (MS)

Digested samples were analysed by liquid chromatography (LC) coupled tandem MS (LC-MS/MS) using an UltiMate® 3000 Rapid Separation LC system (RSLC, Dionex Corporation) coupled to a Q Exactive HF (Thermo Fisher), peptide mixtures were separated using a multistep gradient from 95% A (0.1% formic acid, FA, Thermo Fisher, in water) and 5% B (0.1% FA in acetonitrile) to 7% B at 1 min, 18% B at 58 min, 27% B at 72 min and 60% B at 74 min at 300 nL/min, using a 75 mm × 250 *µ*m, inner diameter 1.7 *µ*m, CSH C18 analytical column (Waters). Peptides were selected for fragmentation automatically by data dependent analysis.

#### Preliminary data analysis using Progenesis QI (Nonlinear Dynamics)

Spectra from multiple samples were automatically aligned using Progenesis QI (version 3.0) with manual placement of vectors where necessary. Peak picking sensitivity was set to 4/5 and features with a charge greater than +4 or fewer than 3 isotopes were excluded from further analysis. In the mouse/human mixed-species experiment, remaining features were searched using Mascot (server version 2.5.1, parser version 2.5.2.0; Matrix Science), against the SwissProt and TREMBL pan-mammalian database for the mixed species dataset or human-only for the stable isotope labelling dataset, or mouse-only for the negative control technical replicate dataset (release-2016_04; non-human and non-mouse identifications were removed). The peptide database was modified to search for alkylated cysteine residues (monoisotopic mass change, +57.021 Da) as a fixed modification, with oxidized methionine (+15.995 Da), hydroxylation of asparagine, aspartic acid, proline or lysine (+15.995 Da) and phosphorylation of serine, tyrosine, threonine (+79.966 Da) as variable modifications. For the PNGase F-treated experiment, the peptide database was modified to search for alkylated cysteine residues (+57.021 Da) as a fixed modification, with oxidized methionine (+15.995 Da), ^18^O-deamidation of asparagine/glutamine (+2.988 Da) and ^16^O-deamidation of asparagine/glutamine (+0.984 Da). A maximum of one missed cleavage was allowed. Peptide tolerance and MS/MS tolerance were set to 8 ppm and 0.015 Da, respectively. Peptide detection intensities were exported from Progenesis QI as Excel (Microsoft) spread sheets for further processing. Deamidated spectrum counts were performed in Scaffold (version 4, Proteome Software) using. DAT files from Mascot, following pre-processing in Progenesis QI as detailed above. We applied quantification calculations to proteins detected with >2 unique (by sequence) peptides. Peptides assigned to proteins with ‘unreviewed’ status in the UniProt database were reassigned to the most abundant ‘reviewed’ protein with sequence identity in the dataset. Peptides shared between different protein groups were excluded from subsequent analysis.

#### Multi-variate regression modelling

To compare different types of regression analysis we used three different types of linear regression algorithms: (A) ordinary least squares (OLS); (B) Linear mixed effects models with Huber residual weights (LME-H); and (C) BayesENproteomics. Further details and references for each method are provided in the following sections. All analysis was performed using code written for Matlab R2015a (The MathWorks Inc). Heatmaps were created using the heatmap.2() function from the gplots R package, using the default options.

#### Imputation of missing values

Monte Carlo Markov Chain (MCMC)-based methods can be used to deduce any unknown value, including missing values in proteomics experiments, provided appropriate conditional distributions can be provided. We used this property to build an adaptive (multiple) imputation (AMI) method that gives consistent accuracy regardless of the type of missingness that dominates a given dataset. We benchmark AMI against other commonly used imputation methods that either exclusively assume values are missing at random (MAR) or missing non-randomly (MNR). MAR imputation was performed using K-Nearest Neighbours (KNN) – where K = 11 or Bayesian principal component analysis (BPCA). KNN, which fills missing values using the average of the K nearest (by Euclidian distance) peptides, was performed using the knnimpute() Matlab function. BPCA was selected as a representative example of PCA-based methods that decompose the dataset and reconstruct the missing values using the principal components. BPCA imputation was performed using Matlab code available from http://ishiilab.jp/member/oba/tools/BPCAFill.html (Oba et al., 2003). MNR imputation was performed by random sampling from a down-shifted Gaussian distribution (DGD) as described by Tyanova et al. (2016) (albeit for protein-level imputation). Briefly, for each peptide, minimum non-missing peptide intensities were averaged to model the centre of a normal distribution with a width equal to 0.3 times that peptide’s σ (standard deviation). The mean (i.e. centre) of this normal distribution was negatively shifted by 1.6σ to model low abundance missingness or “missing non-randomly” (MNR), Missing values were then randomly imputed from this downshifted normal distribution. As an example of model-based imputation, DanteR (Karpievitch et al., 2009), downloaded from https://omics.pnl.gov/software/danter was run in RStudio (version 1.1.453) running R 3.5.2.

AMI uses a logistic regression to determine whether missing values are MAR or MNR and imputes from appropriate conditional distributions. Logistic regression, similar to that employed by Li et al. (2011), was used to discern whether a given missing value was MAR or MNR as in (3).

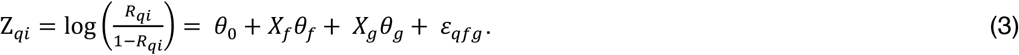

Where *R*_*q*_ represents a binary vector for protein *q* with elements *r*_1_, … *r*_*n*_ denoting whether an observation was missing (*r*_*i*_ = 1) or not (*r*_*i*_ = 0) and Z_*q*_ represents the logit transform of R_*q*_to enable estimation of regressor coefficients *θ* using linear regression. As Z_*qi*_ ∈ {−∞, +∞}, we set minimum and maximum values for Z_*q*_ to −10 (corresponding to a probability for observation *i* being missing, *r*_*i*_ < 0.00005) and 10 (*r*_*i*_ > 0.99995). *X*_*f*_ and *X*_*g*_ represent binary design matrices denoting the peptide (*f*) and treatment group (*g*) from which a given observation is derived. *θ*_*j*_ (*j* = 1…, (*p*_*f*_ + *p*_*g*_ + 1)) represents a 1 × (*p*_*f*_ + *p*_*g*_ + 1) vector of regressor coefficients; *p*_*f*_ and *p*_*g*_ represent the number of elements in *θ*_*f*_ and *θ*_*g*_, respectively. *θ*_*f*_ and *θ*_*g*_ denote whether observations from a given peptide or treatment correlate with values in *R*_*q*_ (i.e. whether probability of “missingness” increases when looking at intensities from particular peptides or from particular experimental treatment groups). The intercept term, *θ*_0_ denotes the intrinsic probability of missingness for that protein. If *θ*_*j*_ > *θ*_0_ and *θ*_*j*_ > 0 we inferred that missing observations associated with *θ*_*j*_ were MNR, and MAR otherwise. MAR and MNR missing values were imputed as part of each Gibbs sampler iteration as in (4) and (5).

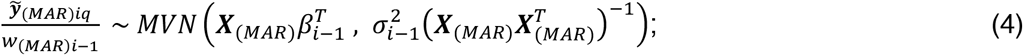

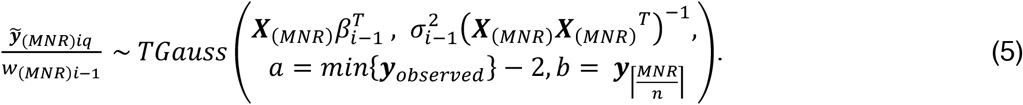

Where 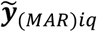 represents a vector of score-weighted (see below) MAR log_2_(intensity) values for protein *q* at the *i*^*th*^ iteration of the Gibbs sampler. 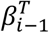 is the vector of parameter estimates and 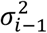 is the residual variance from the *i-1*^*th*^ iteration of the Gibbs sampler, respectively. Missing values in 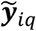 are sampled according to the multivariate normal distribution (MVN) described in (4) (see also section 3.1 in Zeng et al. (2017)). MNR missing values were imputed from a truncated Gaussian (TGauss) with an upper limit determined by the percentile of observed log_2_ intensity values corresponding to the fraction of values that are deemed MNR 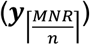 from the missingness regression model in (3). In (5), *a* and *b* represent the lower and upper limits of the TGauss distribution. Thus, if 5% of observations for a given protein are determined to be MNR according to (3), then the cut-off for the truncated Gaussian is equal to the 5^th^ percentile of observed log_2_ intensity values. New missing values were imputed for each iteration of the Gibbs sampler. The multiple imputation strategy implemented in BayesENproteomics thus accounted for the inherent uncertainty in imputation by basing the resulting *β* estimates (and subsequent hypothesis testing) on distributional estimates of missing values (Schafer and Olsen, 1998) rather than fixed point estimates as in OLS and LME-H, where single random samples could strongly influence individual protein fold change estimates.

#### Weighting of residuals based on confidence of peptide identification

Identification of PTM’d peptides was performed by the inclusion of variable modifications during peptide database searching. Inclusion of multiple variable modifications was found to increase the number of false positive peptide identifications. The number of false-positive identifications was reduced by discarding peptides with low Mascot scores using a standard FDR cut-offs based on identification p-values. We employ a Benjamini-Hochberg FDR (Benjamini and Hochberg, 1995) cut-off of < 0.2.

BayesENproteomics also employed a novel heuristic outlier weighting scheme that weighted against outlier peptides (which may possess biologically relevant PTMs), particularly if confidence in their identification was low. In this case we used Mascot scores as our indicator for peptide identification confidence. Firstly, Mascot scores, *S*_1_…, *S*_*n*_ were scaled by dividing them by a modified Bonferroni-like cut-off (similar to that described on the Mascot website, http://www.matrixscience.com/help/interpretation_help.html, accessed 23/10/17) and adjusted so that all the highest scoring peptides were weighted equally, as in (6) with values between 0 and 1.

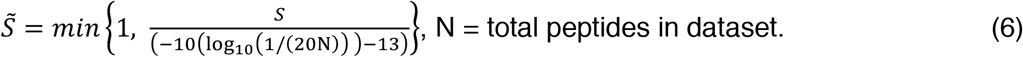

During each iteration of the Gibbs sampler, an n-dimensional vector of weights, *w* with elements *w*_1_…, *w*_*n*_, was calculated using a variation of the automatic outlier detection and weighting method in (Ting et al., 2007). Initially, there was no *a priori* reason to exclude - or diminish the influence of - peptides that have passed the initial FDR screen. Instead, we opted to weight in favour of those peptides that either have high Mascot scores or low residuals (7). Transformed Mascot scores in 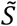 were used to parameterize a binomial distribution giving *Ŝ* that would determine if observations from that were favourably weighted each Gibbs sampler iteration (8).

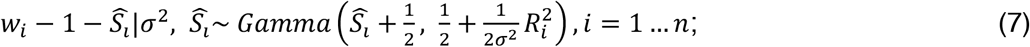

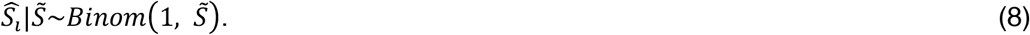

Where ***R*** is a vector of residuals with elements *R*_1_ … *R*_*n*_, *R* = ***y*** − ***X**β*. Observations were then weighted by multiplying each row of ***X*** and each value of ***y*** by their respective weight calculated in (7), (9) and (10).

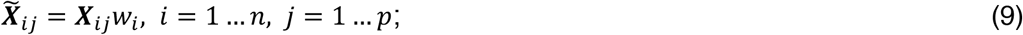

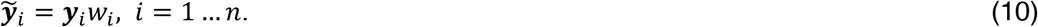

Where 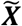 and 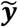 are the weighted design matrix and response vector, respectively.

#### Linear regression implementation

Peptide-based linear regression modelling has previously been shown to possess greater statistical power and accuracy than summarization models when detecting differentially abundant proteins (Goeminne et al., 2015). Here we compare three different models for calculating differentially abundant proteins and PTMs:

##### (A) Ordinary least squares (OLS)

Differential protein abundance was calculated using the simple linear model (Choi et al., 2014; Clough et al., 2009) shown in (1), using the fitlm Matlab function. For calculating differential PTM abundance, the log_2_ fold change for each PTM’d peptide was normalised to the log_2_ fold change calculated for parent protein abundance. In cases where a single PTM site was shared by 2 or more peptides (i.e. missed cleavages), the most abundant one was used.

##### (B) Linear mixed-effects models with Huber residual weights (LME-H)

The ridge regression/mixed-effects algorithm developed by Goeminne et al. (2016), wherein peptides were modelled as random effects (i.e. they were assumed to be randomly sampled from a larger population and that they accurately modelled the variance of that population) using the more complex model shown in equation (2). Goeminne et al. (2016) exploited the link between ridge regression and mixed effects models to assign each *β*_*j*_ a specific penalty, *λ*_*j*_ (where 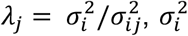 = residual variance of protein *i*, 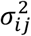 = variance of coefficient *β*_*j*_ for protein *i*, *j* = 1, 2 … *p*). We recapitulated this algorithm using the fitglme Matlab function, including Huber weighting of residuals. To calculate changes in PTM abundances, peptide:group and peptide:donor interaction effects (*β*_*f*:*g*_, and *β*_*f*:*d*_ in (2), respectively) were included as random effects with the size of the resulting interaction *β*s denoting changes in the abundance of that peptide in response to treatment or donor effects, respectively.

##### (C) BayesENproteomics

This method utilises a novel Bayesian linear regression algorithm with elastic net Regularisation based on the hierarchical model detailed by Kyung et al. (2010). Bayesian methods employ Regularisation based on the prior distribution parameters were estimated from, with elastic net Regularisation being equivalent to sampling from an intermediate Gaussian/Laplacian prior. Sampling was performed using a Gibbs sampler with the maximum number of iterations for each protein set at 25 × *runs* + *p*_*f*_ (*runs* = total number of MS runs to be analysed, *p*_*f*_ = number of peptides), or 1000 – whichever was higher – with half of these as burn-in iterations. The full hierarchical model for a single protein is detailed in equations (11)–(15).

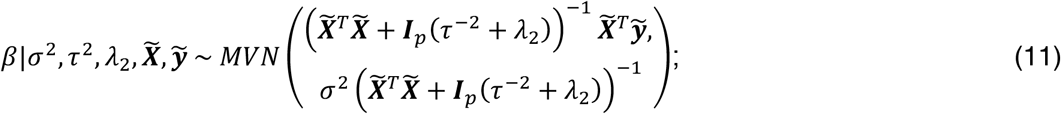

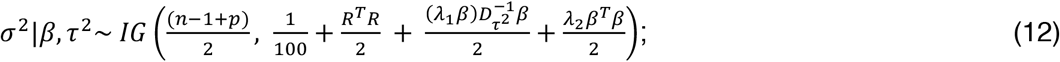

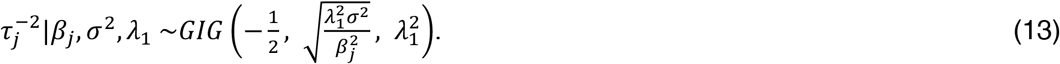

Let *τ*^−2^ represent a vector of latent variables 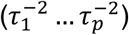 – sampled from the generalised inverse Gaussian (GIG) distribution in (13) using the sampler in Makalic and Schmidt (2016) – for each *β*_*j*_ such that larger values of 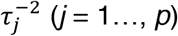 result in *β*_*j*_ being shrunk towards zero. 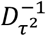 denotes a diagonal matrix with elements 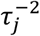, *j* = 1…, *p*. Residual variance (*σ*^2^) is sampled from an inverse gamma (IG) distribution in (11). To enforce sparsity, we employ two Regularisation hyperparameters, *λ*_1_ and *λ*_2_, with different conditional distributions, specifying LASSO (14) and ridge hyperparameters (15), respectively. Notably, while overall covariance is controlled by *λ*_1_ through its effect on 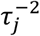, each *β*_*j*_ is given its own L_2_ Regularisation hyperparameter, 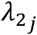, similar to the LASSO-like “horseshoe” estimator (Makalic and Schmidt, 2016). This leads to smaller coefficients (i.e. “noise”) being more aggressively shrunk towards zero compared to larger coefficients (i.e. “signal”), compared to regression using scalar Regularisation hyperparameters that lead to constant shrinkage across all *β*s. 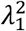 and *λ*_2_ are sampled from gamma distributions of the form *Gamma*(*a*, *b*), with a posterior mean of 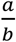.

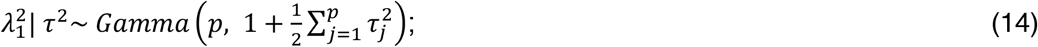

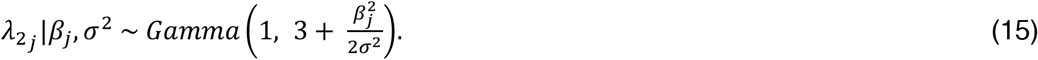

Estimates of parameters were taken as means of Gibbs-sampled posterior distributions made from post-burn-in iterations.

### Pathway analysis using linear modelling (PALM)

PALM using Reactome (Fabregat et al., 2018; Milacic et al., 2012) pathway annotations was performed as described above using logged fold changes (calculated from the three methods detailed above) as the response variable (***y***) according to the model for a given Reactome pathway:

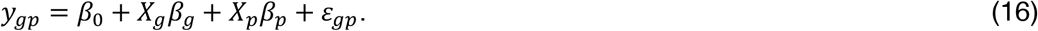

Where *β*_*g*_ and *β*_*p*_ denote pathway-level effect sizes due to experimental treatment *g* and protein *p*. Residual weights (*w*) were set to 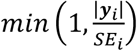, where *SE*_*i*_(*i* = 1… *n*) is the standard error of that protein fold change estimate, ***y***_*i*_. The code allows the user to limit estimation to only those pathways with a minimum observed number of proteins present in a given dataset; here, we set the minimum to be five. Comparisons to PALM were performed using Reactome’s own service (https://reactome.org) or PANTHER (http://pantherdb.org; Mi et al., 2017) using the enrichment option with the default settings.

#### Empirical Bayes correction of protein variances

All methods included Empirical Bayes correction for stabilising variance estimation (Kammers et al., 2015; Smyth, 2004). Empirical Bayes-modified t-tests were used to calculate p-values for determining significance of differentially abundant proteins and PTMs.

#### Data availability

All quantification was performed on Progenesis QI peptide-level outputs using code written for Matlab R2015a (The MathWorks Inc). Progenesis QI peptide ion outputs used in this study, along with protein/pathway-level output files as. csv are freely available from www.github.com/VenkMallikarjun/BayesENproteomics. BayesENproteomics code is available for Matlab (used in this manuscript; www.github.com/VenkMallikarjun/BayesENproteomics) and Python3 (www.github.com/VenkMallikarjun/BENPPy). The raw data have been deposited to the ProteomeXchange with identifiers PXD012784 (mouse skin negative control), PXD012782 (PNGase F stable isotope labelling PTM quantification benchmark) and PXD012772 (human:mouse mixed species protein quantification benchmark).

## Acknowledgements

VM was supported by a studentship from the Sir Richard Stapley Educational Trust. JS was funded by a Biotechnology and Biological Sciences Research Council (BBSRC) David Phillips Fellowship (BB/L024551/1). Mass spectrometry was carried out at the Wellcome Centre for Cell-Matrix Research (WCCMR; 203128/Z/16/Z) Biological Mass Spectrometry Core Research Facility. The authors thank: Prof. Tim Board (Wrightington Hospital) for provision of tissue samples for MSC isolation; Prof. Matthew Hardman and Dr Charis Saville for provision of mouse tissue; Drs Ronan O’Cualain, Stacey Warwood and David Knight for Core Facility support; and Dr Craig Lawless, Prof. Ilaria Bellantuono and Prof. Simon Hubbard for constructive criticism of the manuscript.

## Author contributions

VM designed and implemented the algorithm and developed the benchmark datasets used in this manuscript. VM, SMR and JS wrote the paper.

