## Supplementary Figure 1 for "BayesENproteomics: Bayesian elastic nets for quantification of proteoforms in complex samples"

*Mallikarjun et al.*

### Supplementary Figure 1

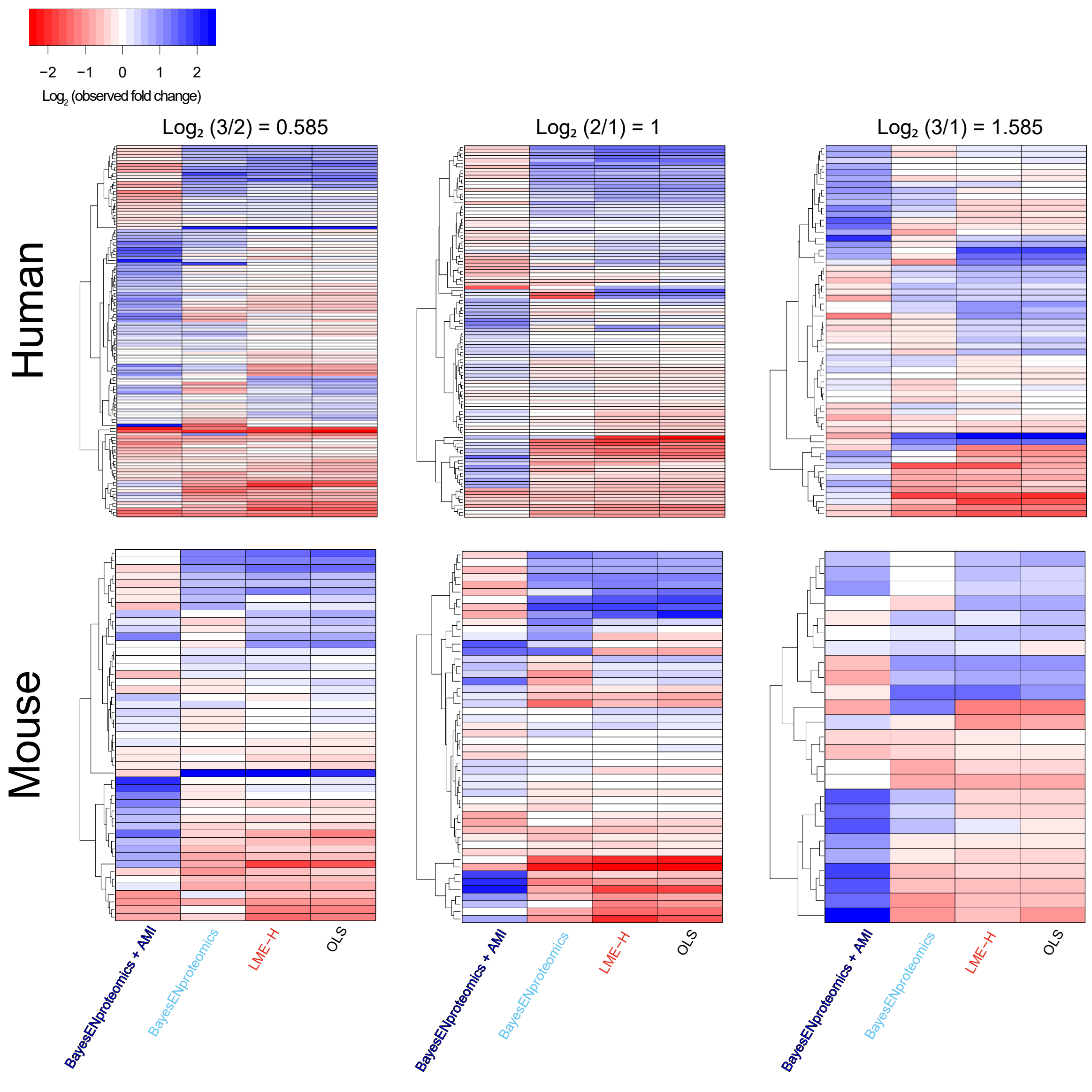

**Figure S1. Consistently incorrect fold change estimates mostly determined by imputation method, rather than regression method.**

Heatmaps showing fold change estimates for those proteins that presented an incorrect (i.e. $< 0$) fold change in any of the mixed species comparisons under any of the regression methods in Figure 3. While BayesENproteomics worsened some and improved others, the most notable differences were caused by changing imputation method from DGD to AMI, with DGD being optimal (i.e. resulting in a correct, positive $\log_2$ fold change, blue) for some while AMI appeared optimal for others.
